# eQTL regulating Transcript Levels Associated with Diverse Biological Processes in Tomato

**DOI:** 10.1101/040592

**Authors:** Aashish Ranjan, Jessica M. Budke, Steven D. Rowland, Daniel H. Chitwood, Ravi Kumar, Leonela Carriedo, Yasunori Ichihashi, Kristina Zumstein, Julin N. Maloof, Neelima R. Sinha

## Abstract

Variation in gene expression, in addition to sequence polymorphisms, is known to influence developmental, physiological and metabolic traits in plants. Genetic mapping populations have facilitated identification of expression Quantitative Trait Loci (eQTL), the genetic determinants of variation in gene expression patterns. We used an introgression population developed from the wild desert-adapted *Solanum pennellii* and domesticated tomato *Solanum lycopersicum* to identify the genetic basis of transcript level variation. We established the effect of each introgression on the transcriptome, and identified ~7,200 eQTL regulating the steady state transcript levels of 5,300 genes. Barnes-Hut *t*-distributed stochastic neighbor embedding clustering identified 42 modules revealing novel associations between transcript level patterns and biological processes. The results showed a complex genetic architecture of global transcript abundance pattern in tomato. Several genetic hotspots regulating a large number of transcript level patterns relating to diverse biological processes such as plant defense and photosynthesis were identified. Important eQTL regulating transcript level patterns were related to leaf number and complexity, and hypocotyl length. Genes associated with leaf development showed an inverse correlation with photosynthetic gene expression but eQTL regulating genes associated with leaf development and photosynthesis were dispersed across the genome. This comprehensive expression QTL analysis details the influence of these loci on plant phenotypes, and will be a valuable community resource for investigations on the genetic effects of eQTL on phenotypic traits in tomato.

## Introduction

The genetic basis of many qualitative and quantitative phenotypic differences in plants has been associated with sequence polymorphisms and the corresponding changes in gene function. However, differences in the levels of steady state transcripts, without underlying changes in coding sequences, also significantly influence plant phenotypes. Closely related plant species often have little coding sequence divergence, nonetheless the related species often develop unique physiological, metabolic, and developmental characteristics indicating that patterns of gene expression are important in species-level phenotypic variation (Kliebenstein, 2009; Koenig et al., 2013). Phenotypic differences attributed to variations in gene expression patterns have been found to influence disease resistance, insect resistance, phosphate sensing, flowering time, circadian rhythm, and plant development (Kroymann et al., 2003; Werner et al., 2005; Clark et al., 2006; Zhang et al., 2006; Svistoonoff et al., 2007; Chen et al., 2010; Hammond et al., 2011).

Global transcript level changes across precise genetic backgrounds have been used to define expression Quantitative Trait Loci (eQTL) by identifying genomic regions responsible for the variation in transcript levels (Jansen and Nap, 2001; Kliebenstein, 2009; Druka et al., 2010; Chitwood and Sinha, 2013). An eQTL is a chromosomal region that drives variation in gene expression patterns (i.e., steady state transcript abundance) between individuals of a genetic mapping population and can be treated as a heritable quantitative trait (Brem et al., 2002; Kliebenstein, 2009; Cubillos et al., 2012). Depending upon the proximity to the gene being regulated, eQTL can be classified into two groups: *cis*-eQTL when the physical location of an eQTL coincides with the location of the regulated gene, and *trans*-eQTL when an eQTL is located at a different position from the gene being regulated (Kliebenstein, 2009). eQTL studies with the model plant Arabidopsis showed that *cis*-eQTL have a significant effect on local expression levels, whereas *trans*-eQTL often have global influences on gene regulation (DeCook et al., 2006; West et al., 2007; Holloway and Li, 2010). Global eQTL studies also identified *trans*-acting eQTL hotspots, which contain master regulators controlling the expression of a suite of genes that act in the same biological process or pathway. For example, eQTL hotspots in Arabidopsis co-locate with the *ERECTA* locus, which has been shown to pleiotropically influence many traits including those regulating morphology (Keurentjes et al., 2007). Similarly the rice *sub1* locus, which regulates submergence tolerance by controlling internode and leaf elongation, controls the activity of ethylene response factors with significant *trans* effects (Fukao et al., 2006). In addition, the eQTL identified using pathogen challenged tissues in barley were enriched for genes related to pathogen response (Chen et al., 2010; Druka et al., 2010). Thus eQTL analyses have the potential to reveal a genome-wide view of the complex genetic architecture of gene expression regulation, the underlying gene regulatory networks, and may also identify master transcriptional regulators.

Cultivated tomatoes, along with their wild-relatives, harbor broad genetic diversity and large phenotypic variability (Moyle, 2008; Ranjan et al., 2012). Wide interspecific crosses bring together divergent genomes and hybridization of such diverse genotypes leads to extensive gene expression alterations compared to either parent. Introgression lines (ILs), developed by crosses between wild-relatives and the cultivated tomato to bring discrete wild-relative genomic segments into the cultivated background, have proved to be a useful genetic resource for genomics and molecular breeding studies. These ILs may vary in the size of introgresssed region that may range from a few genes to more than a thousand genes. ILs developed from the wild desert-adapted species *Solanum pennellii* and domesticated *Solanum lycopersicum* cv. M82 have proved to be a useful genetic resource (Eshed and Zamir, 1995; Liu and Zamir, 1999). This population has been successfully used to map numerous QTL for metabolites, enzymatic activity, yield, fitness traits, and developmental features, such as leaf shape, size, and complexity (Frary et al., 2000; Holtan and Hake, 2003; Fridman et al., 2004; Chitwood et al., 2013; Muir et al., 2014). Comparative transcriptomics for the two parents enabled identification of transcript abundance variation potentially underlying trait differences between species (Koenig et al., 2013). However, the genetic regulators of these transcriptional differences between the species still need to be elucidated. Therefore, we used a genomics approach in combination with statistical methods to identify the genetic basis of transcript level variation in tomato using the *S. pennellii* introgression lines.

Here we report on a comprehensive transcriptome profile of the ILs, a comparison between the transcript abundance patterns of the ILs and the cultivated M82 background (differential gene expression – DE), as well as a global eQTL analysis to identify patterns of genetic regulation of transcript abundance in the tomato shoot apex. We have identified more than 7,200 *cis*-and *trans*-eQTL in total, which regulate the transcript abundance of approximately 5,300 genes in tomato. Additional analyses using Barnes-Hut *t*-distributed stochastic neighbor embedding (BH-SNE) (van der Maaten, 2013) identified 42 modules revealing novel associations between transcript abundance patterns and biological processes. The transcript abundance patterns under strong genetic regulation are related to plant defense, photosynthesis, and plant developmental traits. We also report important eQTL regulating steady state transcript abundance pattern associated with leaf number, complexity, and hypocotyl length phenotypes.

## Results and Discussion

### Transcriptome Profiling and Global eQTL analysis

RNAseq reads obtained from the tomato shoot apex with developing leaves and hypocotyl were used to identify differentially expressed (DE) genes at the transcript level between each *S. pennellii* IL and the cultivated M82 (Supplemental Dataset 1). The total number of genes differentially expressed for each IL both in *cis* (in this population reflecting “local” level regulation either from within a gene itself or other genes in the introgression) and *trans*, along with the number of genes in the introgression regions, are presented in Figure 1 and Supplemental Table I. There was a strong correlation between the number of genes in the introgression regions and the number of DE genes in *cis* (Supplemental Figure S1A). In contrast, the number of DE genes in *trans* was poorly correlated with introgression size (Supplemental Figure S1B). For example, IL12.1.1, despite having one of the smallest introgressions, showed 96% of ~500 DE genes regulated in *trans* (Supplemental Table I, Supplemental Figure S2). In contrast, IL1.1 and IL12.3, the ILs with highest number of genes in the introgression regions, showed smaller numbers of total and *trans* DE genes (Figure 1, Supplemental Table I, Supplemental Figure S2). These examples suggest that specific loci and not the introgression size determine gene regulation in *trans*. This could, in part, be due to the presence of genes encoding key transcription factors or developmental regulators in the regions with strong influence on transcript expression pattern as is seen in the *ERECTA* containing genomic region in Arabidopsis (Keurentjes et al., 2007). A total of 7,943 unique tomato genes were DE between the ILs and cv. M82, representing approximately one third of the ~21,000 genes with sufficient sequencing depth to allow DE analysis. 2,286 genes, more than one fourth of unique DE genes between the ILs and cv. M82, genes showed transgressive expression patterns, i.e. those genes were differentially expressed at the transcript level for the IL but not for *S. pennellii* compared to cv. M82 (Supplemental Dataset 2 and 3). These data suggest that in addition to protein coding differences, transcriptional regulation of less than one third of all genes accounts for most of the phenotypic and trait differences between the ILs and the cultivated parent.

**Figure 1.**
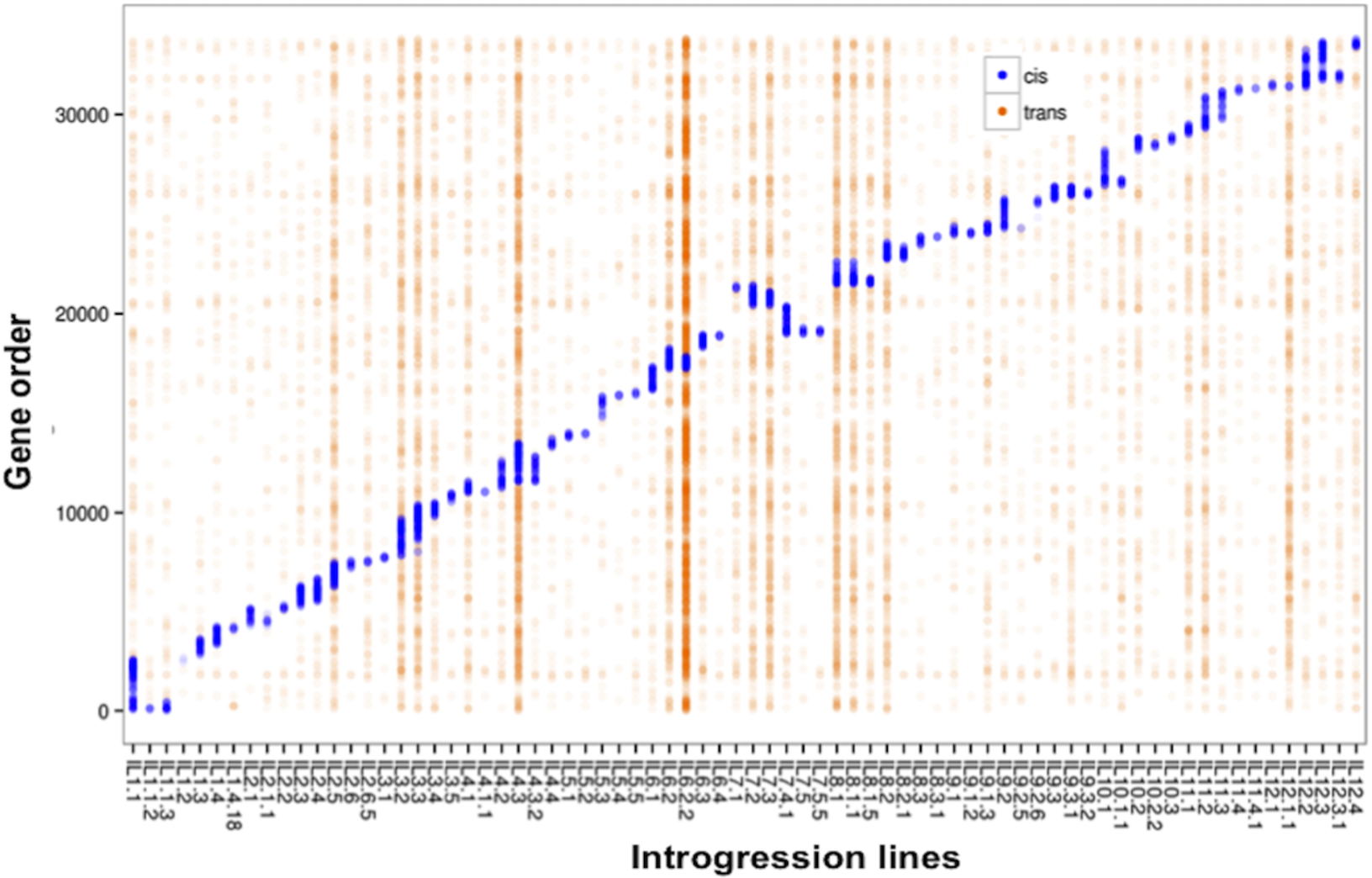
**Transcriptome profile of the tomato introgression lines**. Differentially expressed genes at the transcript level for the ILs compared to cultivated parent M82. Y-axis shows all the tomato genes starting from the first gene on chromosome 1 to the last gene on chromosome 12, and X-axis depicts the individual ILs. Genes differentially expressed within the introgression regions (in *cis*) are shown as blue points and differentially expressed genes in *trans* (outside) the introgression region are shown as orange points.

Identifying eQTL localized to subsets of the introgressions, based on overlaps between them, enabled us to narrow down the regions that contain the regulatory loci. This analysis brings us one step closer to identifying potential candidates that influence transcript abundance patterns in tomato. We identified 7,225 significant eQTL (bins) involving 5,289 unique genes across the 74 ILs (Figure 2; Supplemental Dataset 4). These 7,225 significant eQTL (located in bins) were designated as *cis, trans*, or *chromo0* (unmapped transcripts) as defined in the methods and illustrated in Supplemental Figure S3, and either up or down based in increase or decrease in transcript levels. This correlation resulted in a total of 1,759 *cis*-up and 1,747 *cis*-down eQTL, 2,710 *trans*-up and 920 *trans*-down eQTL, and 51 *chromo0*-up and 38 *chromo0*-down eQTL (Spearman’s rho values, Supplemental Figure S4, Supplemental Table II). The majority of genes (over 4,000 out of 5,289) are under the regulation of a single eQTL (3,134 *cis*, 1,014 *trans*, and 19 *Chromo0*; Supplemental Figure S5). This observation shows the predominance of *cis*-eQTL for genetic regulation of transcript abundance in the tomato ILs. Similar correlation between transcript-level variation and genome-wide sequence divergence within seven Arabidopsis accessions was reported to be due to *cis* control of a majority of the detected variation (Kliebenstein et al., 2006).

**Figure 2.**
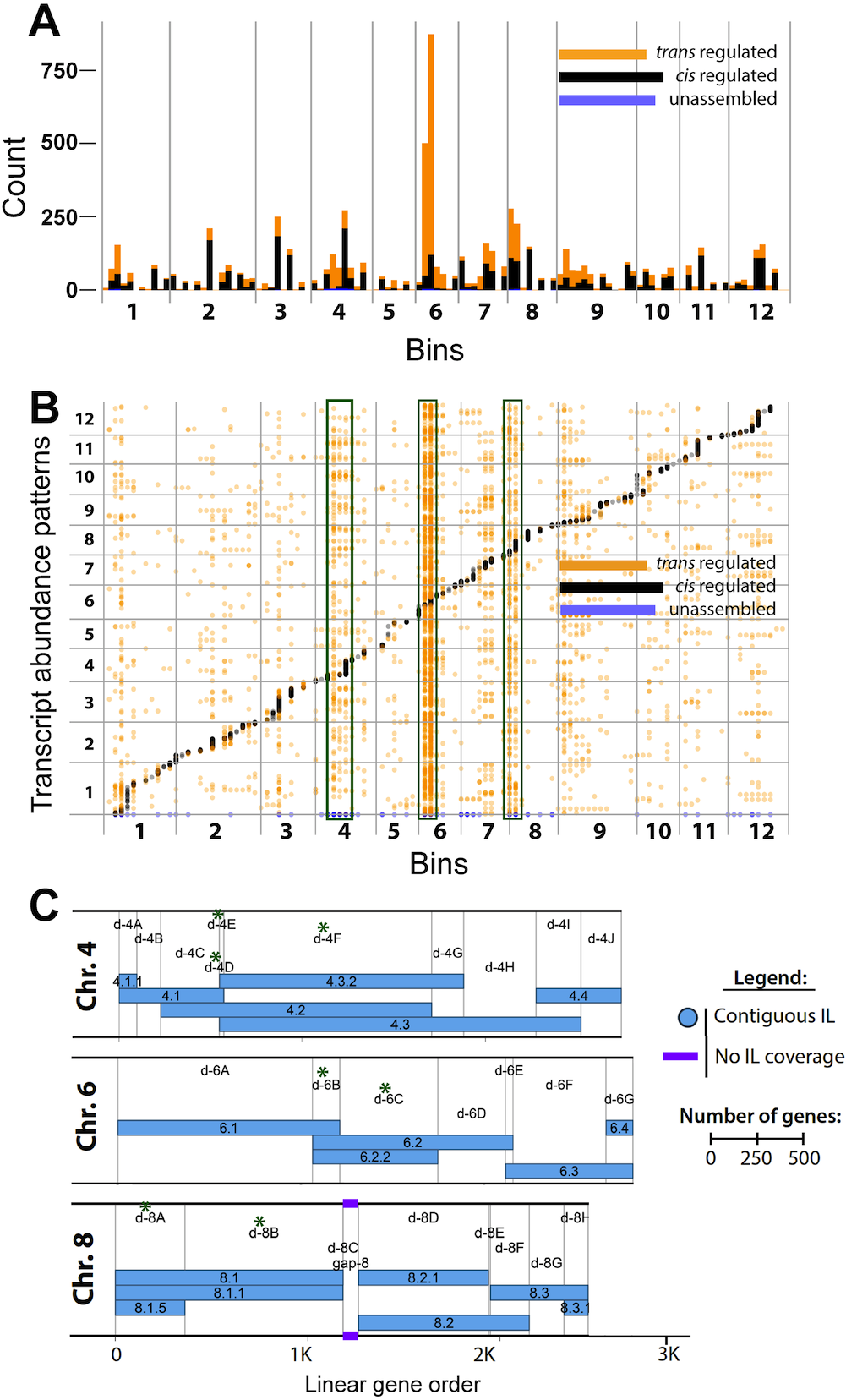
***Cis*-and *Trans*-eQTL plotted by bin across the 12 chromosomes of *S. lycopersicum* cv. M82**. A) Stacked bar graph showing the sum of the number of eQTL mapping to each bin. B) Dotplot showing each eQTL arranged vertically by bin and horizontally by the location of the transcript abundance pattern it regulates. Bins with the largest numbers of *trans*-eQTL (4D, 4E, 4F, 6B, 6C, 8A, 8B) are highlighted by green boxes. C) Map of chromosomes 4, 6, and 8 showing the overlapping IL regions, which define the bins (Modified from Chitwood et al., 2013). Bins with the largest numbers of *trans*-eQTL are indicated by green asterisks.

The number of genes regulated by eQTL showed large variation across bins. Bins on chromosomes 6, 8, and 4, such as 6B, 6C, 4D, 8A, and 8B, contain predominantly *trans*-eQTL (Supplemental Dataset 5). In contrast three bins, 1F, 3I, and 8G, that each contains over 100 genes, have no significant *trans*-or *cis*-eQTL and are transcriptionally silent. As expected, bins containing over 100 significant *cis*-eQTLs are scattered across the genome (Supplemental Dataset 5). The abundance of *trans*-eQTL on chromosomes 4, 6, and 8 strengthens the idea of *trans*-eQTL hotspots controlling expression of a large number of transcripts, as reported in other organisms (Brem et al., 2002; Schadt et al., 2003). The resolution in this analysis is at the level of bin, and these significant eQTL likely map to a smaller number of genes within the bins. Functional classification of genes being regulated by these eQTL and phenotypic association with the relevant ILs was undertaken to glean insights into the identity of candidate genes in the bin.

### Clustering eQTL target genes into modules defined by transcript abundance patterns

In order to functionally categorize the eQTL regulated genes, Barnes-Hut *t*-distributed stochastic neighbor embedding (BH-SNE, van der Maaten, 2013) was performed on the 5,289 genes with eQTL to detect novel associations between transcript abundance patterns. This clustering resulted in 42 distinct modules containing 3,592 genes (Figure 3). Seventeen of these modules had significant GO enrichment (p-value <0.05) with each module consisting of transcript abundance patterns either predominately regulated by *cis*-or *trans*-eQTL (Supplemental Table III). To determine which ILs are important for module regulation, the median transcript abundance value of module genes for each IL was calculated and used to identify ILs with significantly altered module steady state transcript level.

**Figure 3.**
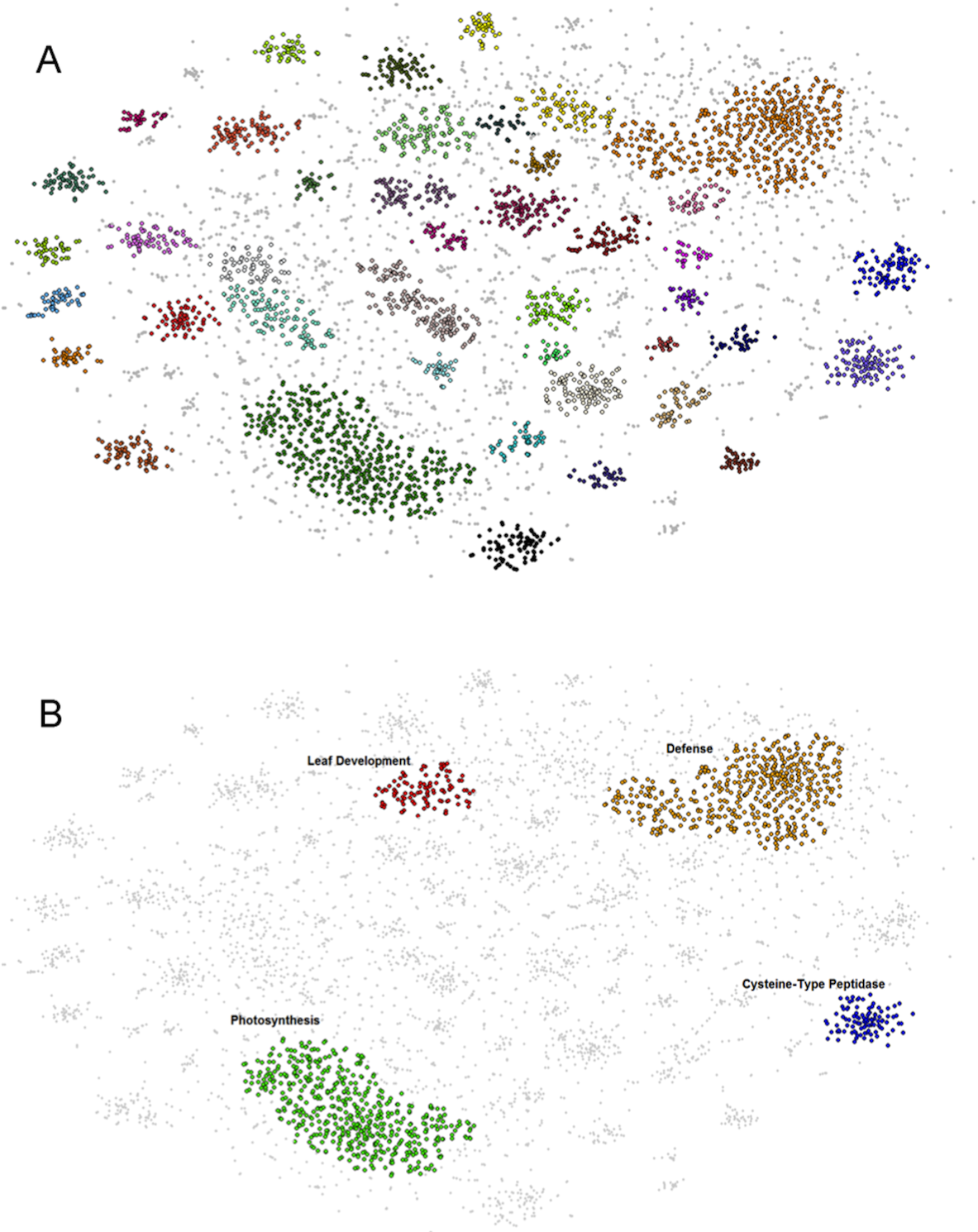
**BH-SNE 2D mapping of eQTL**. A) Forty-two distinct modules identified by DBscan from the mapping generated by BH-SNE analysis. B) The three modules defined as landmark modules: photosynthesis, defense and cysteine-type peptidase activity and the leaf development module’s position within the mapping. Modules are false colored.

Three modules were present in all mappings of the BH-SNE (van der Maaten and Hinton, 2008) determined through iterations of DBscan analysis and GO enrichment, and were designated as landmark modules (Figure 3B; Supplemental Figure S6; Supplemental Dataset 6; Supplemental Table III). The largest module had a GO enrichment for photosynthesis and related processes, and significant *trans*-eQTL scattered widely across the genome with no bin or IL identified as the primary regulating region (Figure 4B; Supplemental Figure S6A; Supplemental Dataset 6 and 7). The second landmark module was enriched for transcript abundance patterns with roles in defense, metabolism, and signaling with the majority of their *trans*-eQTL mapped to IL6.2 and 6.2.2 (Figure 4A; Supplemental Figure S6B; Supplemental Dataset 6 and 8). The third module, which is enriched for transcript abundance patterns with cysteine-type peptidase activity, was predominately composed of genes regulated by *cis*-eQTL on IL 4.2, 4.3, and 4.3.2 (Bins 4E and 4F) (Figure 4C; Supplemental Figure S6C; Supplemental Dataset 6 and 9). A cluster of genes enriched for “peptidase regulation” also emerged from a transcriptome study of leaf development for three tomato species; this cluster was uniquely associated with *S. pennellii* orthologs at the P5 stage of leaf development, indicating that this species has a unique pattern of gene expression, which involves peptidase regulation (Ichihashi et al., 2014) and may be related to leaf maturation and senescence processes (Diaz-Mendoza et al., 2014).

**Figure 4.**
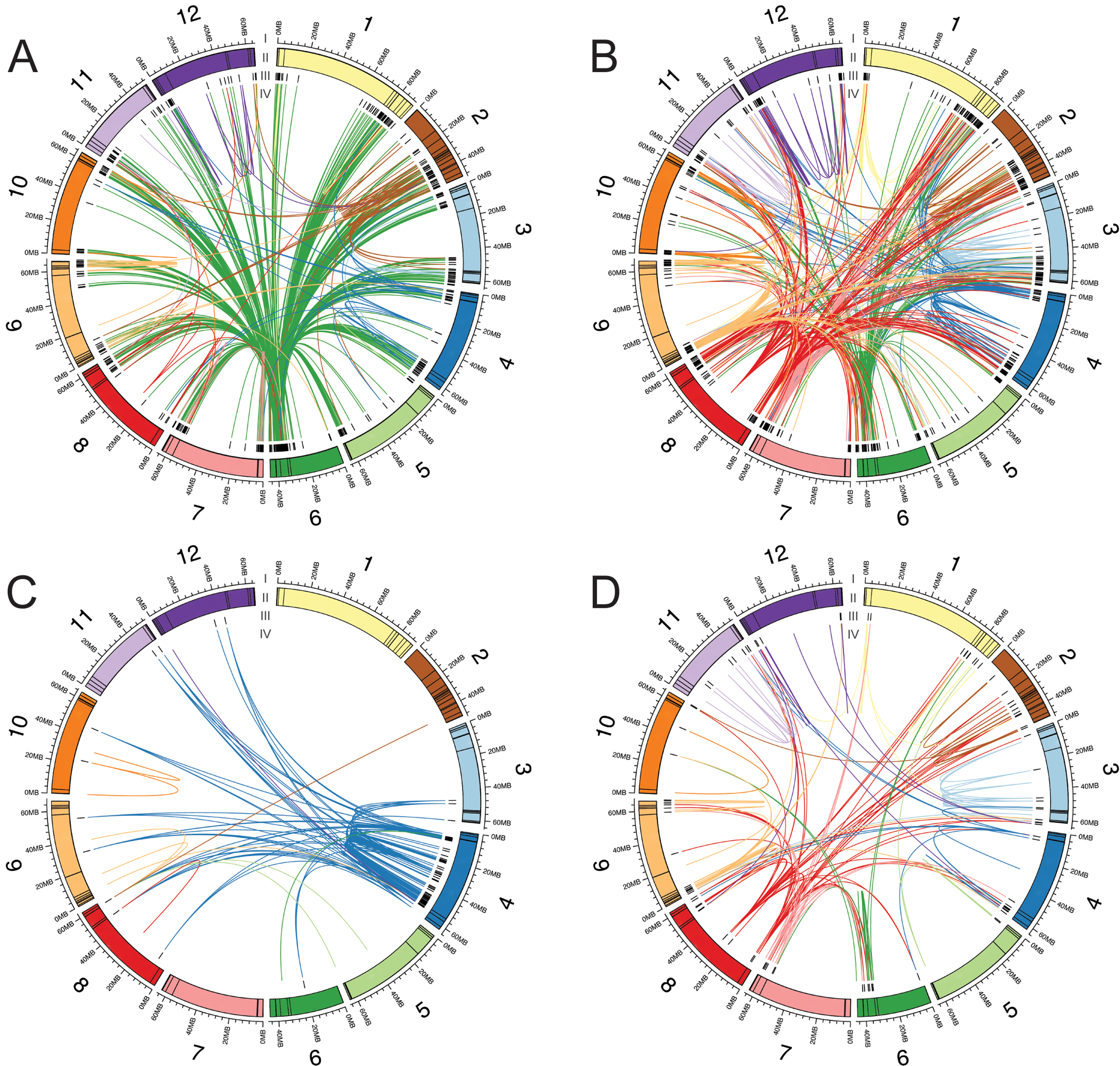
**Connections between eQTL and the genes with correlated transcript level**. Each plot includes the genes with eQTL that were clustered together into a module based on transcript level patterns. A) Defense module. B) Photosynthesis module. C) Cysteine peptidase module. D) Leaf development module. I) The 12 tomato chromosomes in megabases. II) Colored boxes indicate the sizes of each bin. III) Black bars indicate the locations of the genes. IV) Chords connect eQTL to the genes whose transcript level patterns they regulate. Chords are colored by the chromosome location of the eQTL.

### Genetic regulation of transcriptional responses associated with plant defense

One of the landmark modules from the clustering analysis was enriched for transcript abundance patterns related to plant defense (Figure 3B; Supplemental Dataset 8). Therefore we explored the genetic basis of transcriptional changes associated with plant defense. IL6.2 and IL6.2.2, and associated bins 6B and 6C, in particular, influence of the transcriptional responses of genes associated with plant defense and signaling (Supplemental Dataset 1). The genes showing increased steady-state transcript levels in both ILs compared to cv. M82, as well as the genes regulated by the corresponding bins, show enrichment of the GO categories response to stress and stimulus, cell death, defense response, and plant-type hypersensitive response (Supplemental Dataset 10 and 11). Promoter enrichment analysis for these genes showed enrichment of a W-box promoter motif that is recognized by WRKY transcription factors and influences plant defense response (Supplemental Dataset 12 and 13) (Yu et al., 2001). Both the bins, in particular bin 6C, contain genes involved in pathogen, disease, and defense response: such as *NBS-LLR resistance genes, WRKY transcription facfors, Mulfidrug resisfance genes, Penfafricopepfide repeaf-confaining genes, Chifinase*, and *Heaf Shock Profein* coding genes. This transcriptional response in the ILs is also reflected in the morphology of IL6.2.2; the plants are necrotic and dwarfed (http://tgrc.ucdavis.edu/pennelliiils.aspx, Sharlach et al., 2013). Previously, a phenotypic study for the chromosome 6 introgression, specifically a 190Kb region in bin 6C, in a pathogen (*Xanthomonas perforans*)/control experiment was shown to confer hypersensitive response in IL6.2 and 6.2.2 (Sharlach et al., 2013). Taken together, these findings suggest bins 6B and 6C contain master genetic regulators of plant defense response genes, though identification of the causal gene/s that influence many other genes in *trans* will need further genetic dissection of these bins.

### Genetic regulation of transcriptional responses associated with leaf development

Given the striking differences in leaf features between *Solanum pennellii* and cv. M82 that are manifested in many ILs (Chitwood et al., 2013), the IL population provides an excellent system for determining the extent of genetic regulation of genes controlling leaf development. Previous phenotypic and QTL analyses identified many ILs, such as IL4.3, IL8.1.5, IL8.1.1, and IL8.1, harboring loci regulating leaf and plant developmental traits (Holtan and Hake, 2003; Chitwood et al., 2013; Muir et al., 2014). IL4.3, which harbors loci with the largest contribution to leaf shape and shows larger epidermal cell sizes (Chitwood et al., 2013), exhibited decreased steady-state transcript levels for many genes associated with cell division, such as Cyclin-dependent protein kinase regulatorlike protein (CYCA2;3), Cyclin A-like protein (CYCA3;1), and F-box/LRR-repeat protein 2 SKP2A (Supplemental Dataset 1 and 10). In addition, genes showing differences in transcript levels in IL4.3 were enriched for the promoter motifs MSA (M-specific activators that are involved in M-phase specific transcription) and the E2F binding site (Supplemental Dataset 11). Genes with decreased transcript levels in ILs 8.1.5, 8.1.1, and 8.1, also included genes associated with leaf development and morphology, genes encoding WD-40 repeat family protein LEUNIG, Homeobox-leucine zipper protein PROTODERMAL FACTOR 2, and the transcription factor ULTRAPETALA (Supplemental Dataset 1 and 10; Abe et al., 2003; Cnops et al., 2004; Carles et al., 2005).

We, further, investigated the transcript expression dynamics of a set of literature-curated genes related to leaf development (Ichihashi et al., 2014) across the ILs and bins (Supplemental Dataset 14 and 15). A number of canonical leaf developmental genes such as *SHOOT MERISTEMLESS* (Solyc02g081120, *STM*), *GROWTH-REGULATING FACTOR* 1 (*GRF1*, Solyc04g077510), *ARGONAUTE 10* (*AGOI0*, Solyc12g006790), *BELL* (*BEL1*, Solyc08g081400), *LEUNIG* (Solyc05g026480) and *SAWTOOTH 1* (*SAW1*, Solyc04g079830) were differentially expressed at the transcript level in more than five ILs. At the level of bins, genes involved in leaf development were regulated by eQTL scattered widely across the genome (Figure 4D). eQTL(bin)-regulation of leaf developmental genes for some of ILs, such as IL 2.1, 4.3, 5.4, 8.1/8.1.1/8.1.5 and 9.1.2, showing strong leaf phenotypes is summarized in Supplemental Table IV. We then examined the location of literature-curated leaf developmental genes within the identified modules in the BH-SNE mapping (Figure 3). The highest number of literature-curated leaf developmental genes (108) was located in the photosynthesis module, whereas 19 of these genes were located in the leaf development module (Supplemental Figure S7B; Supplemental Dataset 16 and 17), suggesting a relationship between these two modules. Over one third of the transcript expression patterns in the leaf development module have significant eQTL that map to bins 4D, 8A, and 8B (5.4%, 16.2%, and 15.5%, respectively; Supplemental Dataset 18), suggesting that these bins contain important regulators of leaf development. This enrichment of eQTL for specific bins is also consistent with the strong leaf phenotypes for ILs 4.3, 8.1, 8.1.1, and 8.1.5. Altogether DE, eQTL, and BH-SNE results indicate that while there is no obvious master regulatory bin for leaf developmental genes, many are under strong genetic regulation by eQTL distributed throughout the genome (Figure 4D). This observation underscores the highly polygenic regulation of leaf development (Chitwood et al., 2013) as multiple loci, residing in many different chromosomal locations, regulate the expression of key leaf-developmental genes at the transcriptional level.

### Genetic regulation of transcriptional responses associated with photosynthesis

Since photosynthesis GO terms were enriched for the largest module from the clustering analysis (Figure 3B) and there was a correlation between photosynthesis and leaf developmental modules (Supplemental Figure S7B), we examined the genetic regulation of photosynthetic genes by specific ILs and corresponding bins. Genes related to photosynthesis show increased transcript levels across 21 ILs distributed on all chromosomes except chromosome 5 (Supplemental Dataset 10), showing multigenic regulation of photosynthetic traits. Many of these ILs, including 8.1.5, 8.1.1, 8.1 and 4.3, and associated bins showed regulation of genes linked to photosynthesis, chlorophyll biosynthesis, and response to light stimulus (Supplemental Dataset 10 and 11). This observation indicates that ILs may also differ from each other and from the cultivated M82 background in photosynthetic efficiency. However no studies, so far, have investigated the photosynthetic phenotype of these ILs.

To analyze the relationship between the leaf development and photosynthesis modules the median transcript abundance value of all genes in each module was compared resulting in a significant negative correlation (adj r2 = = 0.77; Figure 5). This analysis likely reflects the transition from leaf development to leaf maturation captured in our shoot meristem samples. The genes found in the leaf development module may promote developmental processes such as cell division and maintenance or meristematic potential, whereas the leaf development related genes found in the photosynthesis module may act to suppress this process to allow for maturation of the leaf. The two modules had their most influential eQTL on bins 4D, 8A, and 8B (Supplemental Dataset 6; Supplemental Figure S7A), suggesting that leaf development and photosynthetic genes not only have transcript levels in opposition but also likely share common regulatory loci. This finding is consistent with the link between leaf development and photosynthesis that we established previously by meta-analysis of developmental and metabolic traits (Chitwood et al., 2013).

**Figure 5.**
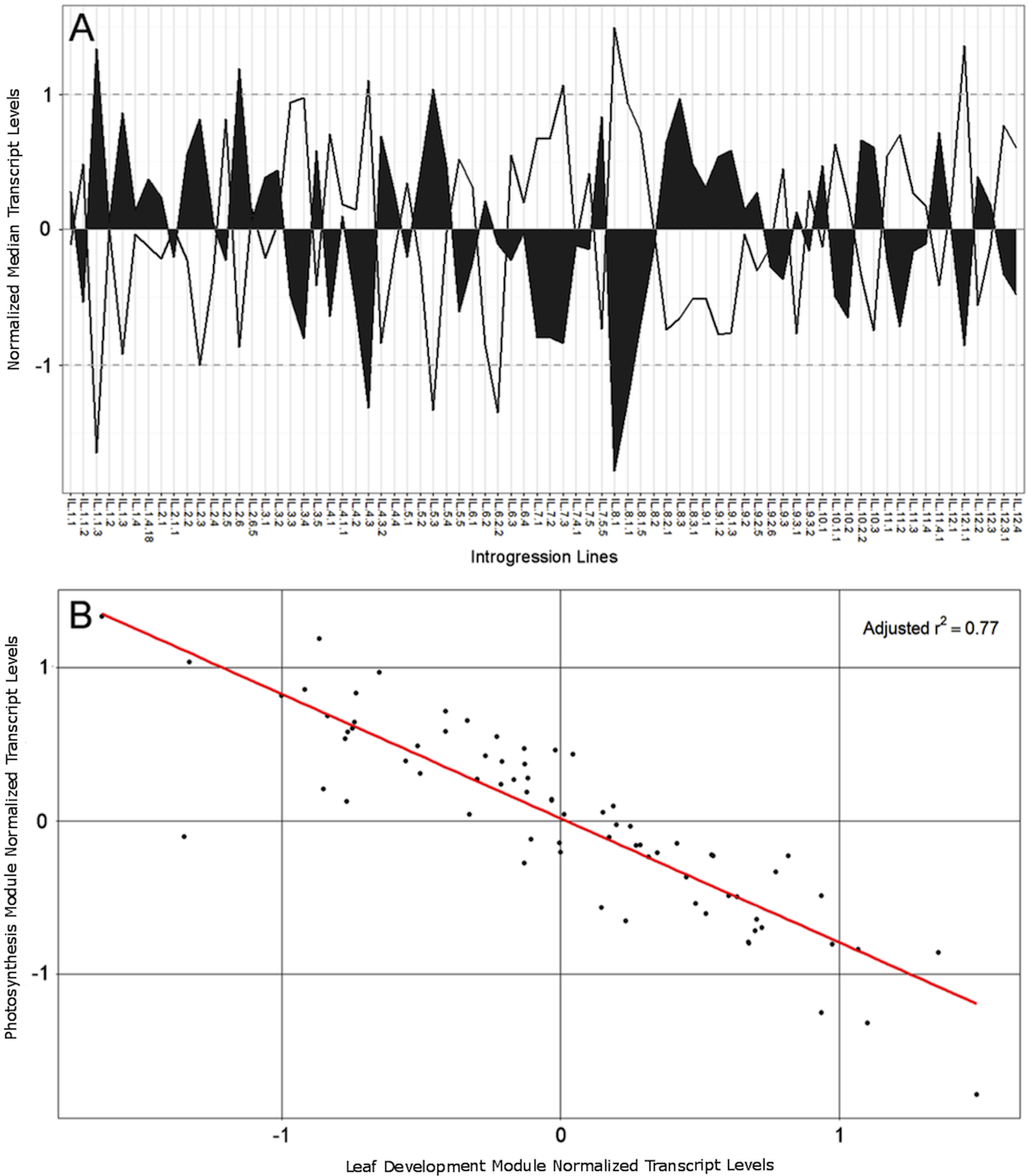
**Median transcript level values for leaf development and photosynthesis related modules and expression correlation**. A) The median transcript level values of a module for each IL are shown. A consistent negative correlation between photosynthesis and Leaf development transcript expression is evident across nearly all 74 ILs. Dashed lines indicate one significant deviation from the module mean transcript level. Filled areas represent the median transcript level of the leaf development module, while open areas indicate the photosynthesis module median transcript levels. B) Leaf development median transcript level versus photosynthesis median transcript level values for each IL show a distinct negative correlation with an adjusted R-squared value of 0.77 (calculated by linear regression in R).

### Dissection of identified eQTL to spatially- and temporally- regulated development

Since the eQTL study used shoot apices that includes the shoot apical meristem (SAM) and developing leaves, we resolved the detected eQTL to specific tissues and temporally-regulated development using previous gene expression data. We analyzed transcript abundance in laser micro-dissected samples representing the shoot apical meristem (SAM)+ + P0 (the incipient leaf) vs. the P1 (the first emerged leaf primordium) that represents transcript levels in the meristem (SAM) and the first differentiated leaf (P1) (Figure 6A). We also analyzed hand dissected samples of the SAM+ + P0-P4 vs. the P5 collected over time (Figure 6B–C), representing genes regulated by vegetative phase change (heteroblasty) (Chitwood et al., 2015).

**Figure 6.**
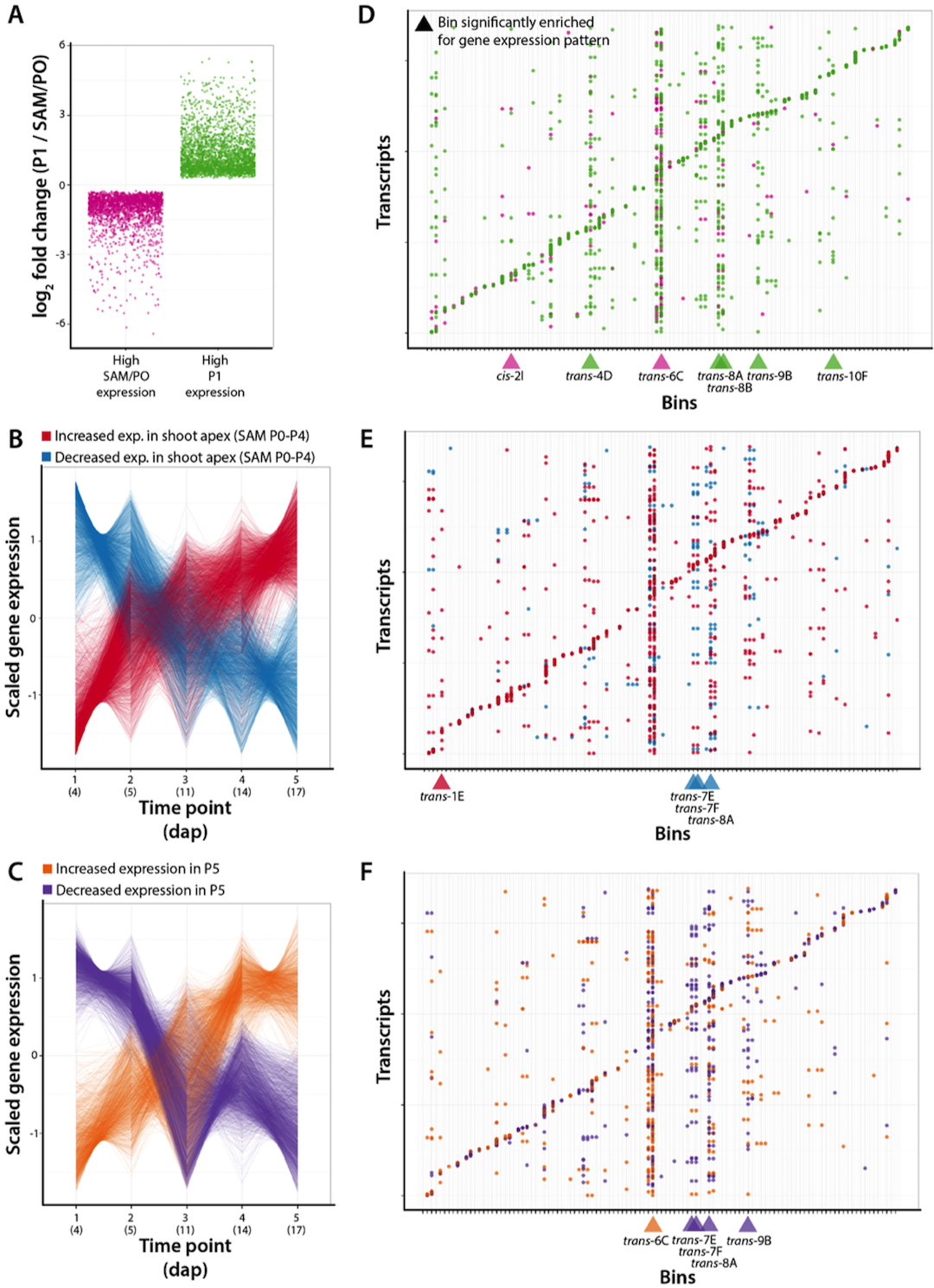
**Enriched gene transcript levels that are controlled by specific bins**. A) Log fold change values (P1/SAM+P0) for previously identified differentially expressed genes with high transcript levels in the SAM + P0 (magenta) vs. P1 (green). B) Scaled transcript level values for previously identified differentially expressed genes with increasing (red) and decreasing (blue) transcript levels over developmental time in the SAM + P0-P4. C) Scaled expression values for previously identified genes with differential levels of transcripts with increasing (orange) and decreasing (purple) transcript levels over developmental time in P5. D) transcripts (y-axis) and bins (x-axis) showing the genetic regulation of transcript abundance (eQTL). Colors indicate SAM + P0 (magenta) and P1 (green) transcripts. Bins enriched for genetically regulating genes with specific transcript expression patterns are indicated below with triangles. E) Same as in D), except showing genes with increasing (red) and decreasing (blue) transcript levels over temporal time in the SAM + P0-P4. F) Same as in D), except showing genes with increasing (orange) and decreasing (purple) transcript levels over temporal time in P5. Previously determined transcript abundance patterns are previously published (Chitwood et al., 2015).

Using a bootstrapping approach, we identified bins statistically enriched for genetically regulating genes with previously identified transcript expression patterns (Figure 6D–F). Except for one instance (cis-regulated genes with high SAM/P0 expression located in bin 2I), bins enriched for transcript expression patterns represented *trans* regulation, hinting at predominant regulation of gene expression patterns mediated by transcription factors at the level of transcription. Most SAM/P0 vs. P1 enriched bins were enriched for P1 transcript expression (Figure 6D). We previously showed that genes with high P1 transcript levels are enriched for photosynthetic-related GO terms, compared to SAM/P0 genes enriched for transcription, cell division, and epigenetics-related GO terms (Chitwood et al., 2015), suggesting a genetic basis at both a functional and tissue-specific level for genes related to photosynthesis expressed preferentially in the P1 compared to the SAM/P0.

Bins enriched for regulation of genes with temporally-dependent steady state transcript levels were mostly associated with genes with decreasing transcript level over time, for both the SAM + P0-P4 and P5 (Figure 6E–F). Interestingly, 3 bins (7E, 7F, and 8A) share enrichment for genes with decreasing transcript levels over time in both the SAM+ + P0-P4 and P5 (Figure 6E–F), suggesting true temporal *trans* regulation, regardless of tissue, by these loci. Broadly, genes with increasing transcript levels over time are associated with transcription and small RNA GO terms in both the SAM + P0-P4 and P5, whereas decreasing transcript levels over time is associated with translation associated GO terms in the SAM + P0-P4 and photosynthetic activity in the P5 (Supplemental Dataset 19).

### Linking leaf and hypocotyl phenotypes to detected eQTL

In order to connect detected eQTL with leaf and hypocotyl phenotypes under two different environmental conditions, we correlated transcript abundance with leaf number, leaf complexity (as measured in Chitwood et al., 2014), and hypocotyl length phenotypes of the ILs grown under simulated sun and shade conditions. Significant correlations with transcript abundance patterns were identified for all three phenotypes analyzed under both treatments (Supplemental Table V). Focusing on a subset of these transcript expression patterns that had associated eQTL enabled us to connect the phenotypes to their regulatory loci (Supplemental Table V).

Genes negatively correlated with leaf number showed enrichment of leaf development GO terms, whereas positively correlated genes showed enrichment of photosynthesis-related GO terms (Supplemental Figure S8A-B; Supplemental Dataset 2 in Chitwood et al., 2014). For the leaf complexity trait, correlations were reversed compared to leaf number (Supplemental Figure S9A-B; Supplemental Dataset 20). The transcript levels of these genes associated with leaf number were predominantly regulated by eQTL on chromosomes 7 and 8 (Supplemental Figure S8C-D), and those of leaf complexity on chromosomes 4, 7, and 8 (Supplemental Figure S9C-D). These results, in combination with DE, eQTL, and BH-SNE, highlight bins on chromosomes 4 and 8 as important genetic regulators of leaf developmental genes.

Five genes were positively correlated with hypocotyl length under simulated shade and one gene (Solyc10g005120) was negatively correlated with hypocotyl length under both sun and shade (Figure 7A; Supplemental Dataset 21). eQTL for the positively correlated genes are located on chromosomes 3, 7, and 11, whereas the single cis-eQTL for the negatively correlated gene, Solyc10g005120 (an uncharacterized *Flavanone 3-hydroxylase-like* gene), was located in bin 10A.1 (Supplemental Figure S10; Figure 7B). The transcript is only expressed in IL 10.1, which has the *S. pennellii* version of the gene and an attenuated shade avoidance response, but is not expressed in IL 10.1.1, which has the M82 version of the gene and a normal shade avoidance response (Supplemental Figure S11). This indicates that genes in bin 10A, the nonoverlapping regions of 10.1 and 10.1.1, are responsible for the shade avoidance response. Bin 10A includes Solyc10g005120, the one gene negatively correlated with hypocotyl length under both sun and shade..

**Figure 7.**
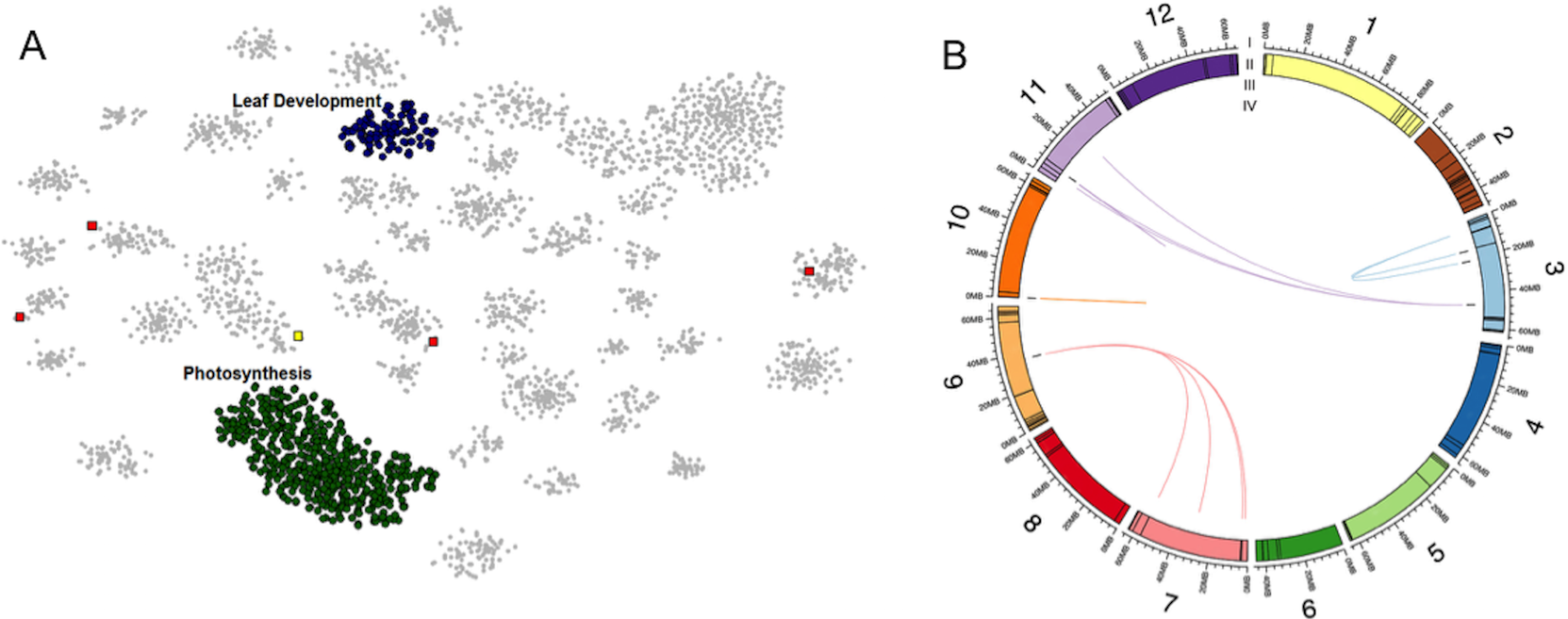
**eQTL regulation of transcript abundance patterns that correlate with hypocotyl length**. A) Forty-two distinct modules identified by DBscan from the eQTL mapping generated by BH-SNE analysis. Modules enriched for genes with leaf development and photosynthesis GO terms are labeled in blue and green, respectively. Genes with transcript levels correlated with hypocotyl length under simulated shade are indicated by squares with positive correlations in red and negative correlations in yellow. B) Genes with transcript levels correlated with hypocotyl length under simulated shade are shown connected to their respective eQTL with chords. I) The 12 tomato chromosomes in megabases. II) Colored boxes indicate the sizes of each bin. III) Black bars indicate the locations of the genes. IV) Chords connect eQTL to the genes whose transcript levels those eQTL regulate. Chords are colored by the chromosome location of the eQTL.

A set of BILs (Backcross Inbred Lines), developed from cv. M82 and *S. pennellii*, provide higher resolution gene mapping with smaller bin sizes (Muller et al., 2016; Fulop et al., 2016). To further explore the role of Solyc10g005120 we used BIL-128, which contains a sub-region of bin 10A and has a secondary introgression on chromosome 2 (Supplemental Figure S10). Influence of the secondary introgression was examined using BIL-033, which-shares the introgression on chromosome 2. BIL-128 has an attenuated shade avoidance response, as does 10.1, whereas BIL-033 undergoes a shade avoidance response similar to that of cv. M82 (Supplemental Figure S11). These results rule out the influence of chromosome 2 genes on the attenuated hypocotyl phenotype and confirm the influence of the bin 10A sub-region, which includes Solyc10g005120, on the attenuated hypocotyl phenotype (Supplemental Figure S10; S11). Solyc10g005120 is an uncharacterized gene and our observations highlight it as a new candidate regulating shade avoidance responses.

## Conclusion

In this study we have investigated the regulation of steady state transcript levels in the progeny of crosses between cultivated tomato (*Solanum lycopersicum* cv. M82) and a wild relative (*Solanum pennellii*). A combination of differential gene expression, eQTL, and clustering analyses provide a comprehensive picture of genetic regulation of transcript expression patterns in this IL population. Our data show that some biological pathways, such as plant defense, are under the regulation of a limited number of loci with strong effects, whereas loci regulating other pathways, such as photosynthesis and leaf development, are scattered throughout the genome most likely with weaker individual effects. We correlated transcript levels with leaf and hypocotyl phenotypes and identified the regulatory regions driving these transcript expression patterns. Coupled with comprehensive phenotyping on these ILs, these data set provide a valuable resource to design strategies to achieve a desirable plant phenotype through genetic manipulation of the transcript abundance of key genes or gene modules. Our ability to predict and understand the downstream effects of genes introgressed from wild relatives on gene expression patterns and ultimately phenotypes will be a critical component of crop plant enhancement.

## Materials and Methods

### Plant Materials, Growth Conditions, and Experimental Design

Plant materials, growth conditions, and experimental design were described in (Chitwood et al., 2013), but are outlined here briefly. Seeds of *Solanum pennellii* ILs (Eshed and Zamir, 1995; Liu and Zamir, 1999) and *Solanum lycopersicum* cv. M82 were obtained either from Dani Zamir (Hebrew University, Rehovot, Israel) or from the Tomato Genetics Resource Center (University of California, Davis). Seeds were stratified in 50% bleach for 2 min., grown in darkness for 3 d for uniform germination before moving to a growth chambers for 5 days. Six seedlings of each genotype were planted per pot for each replicate. The 76 ILs (and two replicates each of cv. M82 and *S. pennellii*) were divided into four cohorts of 20 randomly assigned genotypes. These cohorts were placed across four temporal replicates in a Latin square design as described in (Chitwood et al., 2013). The seedlings were harvested 5 d after transplanting (13 d of growth in total). Cotyledons and mature leaves >1 cm in total length were excluded, and remaining tissues (including the shoot apical meristem) above the midpoint of the hypocotyl were pooled, for all individuals in a pot, into 2-mL microcentrifuge tubes and immediately frozen in liquid nitrogen. Two ILs, IL7.4 and IL12.4.1 were not included in the final analysis due to seed contaminations.

### Growth conditions and quantification of hypocotyl length

Seeds 76 ILs (covering the entire genome) along with the parents were sterilized using 70% ethanol, followed by 50% bleach, and finally rinsed with sterile water. This experiment was replicated three times each in 2011 and 2012. Ten to twelve seeds of each IL were sown into Phytatray II (Sigma-Aldrich) containers with 0.5x Murashige and Skoog minimal salt agar. Trays were randomized and seeds germinated in total darkness at room temperature for 48h. Trays of each IL were randomly assigned to either a sun or shade treatment consisting of 110μMol PAR with a red to far-red ratio of either 1.5 (simulated sun) or 0.5 (simulated shade) at 22°C with 16 hour light / 8 hour dark cycles for 10d. Three genotypes were excluded from the analyses due to poor germination (IL3.3) or their necrotic dwarf phenotypes (IL6.2, 6.2.2). After 10d, seedlings were removed from the agar and placed onto transparency sheets containing a moistened kimwipe to prevent dehydration and scanned using an Epson V700 at 8-bit grayscale at 600 dpi. Image analysis was carried out using the software ImageJ (Abramoff et al., 2004).

For hypocotyl length analysis of backcross inbred lines between *S. pennellii* and *S. lycopersicum* cv. M82, seeds were sterilized in 50% bleach and then rinsed with sterile water. The seeds were then placed in Phytatrays in total dark at room temperature for 72 hours, and then moved to 16 hour light / 8 hour dark for 4d. Seedlings were transferred to soil using a randomized design and assigned to either a sun or shade treatment (as described above) for seven days. Images were taken with a HTC One M8 Dual 4MP camera and hypocotyl lengths measured in ImageJ (Abramoff et al., 2004) using the Simple Neurite Tracer (Longair et al., 2011) plugin.

### RNA-Seq Library Preparation and Preprocessing RNA-Seq Sequence Data

RNAseq libraries were prepared and the reads were preprocessed as described in (Chitwood et al., 2013), and are outlined here. mRNA isolation and RNA-Seq library preparation were performed from 80 samples at a time using a high-throughput RNA-Seq protocol (Kumar et al., 2012). The prepared libraries were sequenced in pools of 12 for replicates 1 and 2 (one lane each) and in pools of 80 for replicates 3 and 4 (seven lanes) at the UC Davis Genome Centre Expression Analysis Core using the HiSeq 2000 platform (Illumina). Preprocessing of reads involved removal of low quality reads (phred score < 20), trimming of low-quality bases from the 39 ends of the reads, and removal of adapter contamination using custom Perl scripts. The quality-filtered reads were sorted into individual libraries based on barcodes and then barcodes were trimmed using custom Perl script.

### Read Mapping and Quantification of Transcript Levels

Mapping and normalization were done on the iPLANT Atmosphere cloud server (Goff et al., 2011). *S. lycopersicum* reads were mapped to 34,727 tomato cDNA sequences predicted from the gene models from the ITAG2.4 genome build (downloadable from ftp://ftp.solgenomics.net/tomato_genome/annotation/ITAG2.4_release/). A pseudo reference list was constructed for *S. pennellii* using the homologous regions between *S. pennellii* scaffolds v.1.9 and *S. lycopersicum* cDNA references above. Using the defined boundaries of ILs, custom R scripts were used to prepare IL-specific references that had the *S. pennellii* sequences in the introgressed region and *S. lycopersicum* sequences outside the introgressed region. The reads were mapped using BWA (Li and Durbin, 2009; Roberts and Pachter, 2013) using default parameters except for the following that were changed: bwa aln: −k 1 −l 25 −e 15 −i 10 and bwa samse: −n 0. The bam alignment files were used as inputs for express software to account for reads mapped to multiple locations (Roberts and Pachter, 2013). The estimated read counts obtained for each gene for each sample from express were treated as raw counts for differential gene expression analysis. The counts were then filtered in R using the Bioconductor package EdgeR version 2.6.10 (Robinson and Oshlack, 2010) such that only genes that had more than two reads per million in at least three of the samples were kept. Normalization of read counts was performed using the trimmed mean of M-values method (Robinson and Oshlack, 2010), and normalized read counts were used to identify genes that are differentially expressed at the transcript level in each IL compared to cv. M82 parent as well as in between two parents, *S. pennellii* and M82. The DE genes for each IL were compared to those between the two parents to identify genes that were differentially expressed for the IL but not for *S. pennellii* compared to cv. M82. Those genes were considered to show transgressive expression pattern at the transcript level for the specific IL, whereas other DE genes were considered to show the transcript expression similar to *S. pennellii*.

### Correlation of phenotype with pattern of steady state transcript levels

Transcript level patterns were correlated with three phenotypes collected from the ILs along with the parents. Normalized estimated read counts with 3-4 independent replicates per IL were log2 transformed prior to the analyses. Leaf number and leaf complexity were collected from the ILs as outlined in Chitwood et al. (2014) under both sun and shade treatments. Hypocotyl lengths were measured as detailed above. To test whether the transcript level for a given gene was correlated with a particular phenotype, boostrapping analyses were performed. transcript levels and phenotype data were randomly permuted (with replacement) using the sample() function against IL and then merged. For each analysis, 1000 replications were performed and the p-values were calculated from the Spearman’s rho value distributions. P-values were adjusted for multiple comparisons using the BH correction (Benjamini & Hochberg, 1995). Significant correlations were identified as those with an adjusted p-value < 0.05 and the mean rho value (the correlation coefficient) was used to designate the correlation as either positive (positive slope) or negative (negative slope). All analyses were implemented using the statistical software R and custom scripts (R Core Team, 2015).

### Methods for eQTL Analyses

eQTL mapping analyses were performed to determine whether the transcript level of a gene is correlated with the presence of a specific introgression from *S. pennellii* into *S. lycopersicum* cv. M82. This correlation was examined at the level of“bin”, with a bin defined as a unique overlapping region between introgressions. Examining eQTL at the bin level enables those eQTL to be mapped to considerably smaller intervals than the ILs themselves (Liu and Zamir, 1999). eQTL mapping analyses were performed on the normalized estimated read counts with 3-4 independent replicates per IL, which were log2 transformed prior to the analyses. To test whether the transcript level for a given gene is correlated with the presence of a particular bin, a Spearman’s rank correlation test was used with ties resolved using the midrank method. P-values were adjusted for multiple comparisons using the BH correction (Benjamini and Hochberg, 1995). Significant eQTL were identified as those with an adjusted p-value < 0.05 and Spearman’s rho (the correlation coefficient) was used to designate the eQTL as up (positive slope) or down (negative slope). Significant eQTL were also designated as *cis* (defined as local gene regulation within the same bin) – if the gene was located on the bin with which it is correlated, *trans* (distant) – if the gene was correlated with a bin that is neither the bin it is on nor a bin that shares an overlapping IL with the correlated bin, or *Chromo0* – if the gene lies in the unassembled part of the genome. When a gene has a designation *cis*-eQTL, and a secondary correlation was found with a bin that shares an overlapping introgression, this secondary correlation was not designated as an eQTL. When a gene does not have a designated *cis*-eQTL and a correlation was found with a bin that shares an overlapping introgression, this correlation was designated as a *trans*-eQTL. All analyses were implemented using the statistical software R and custom scripts (R Core Team, 2015).

### Methods for eQTL clustering analysis

**Data Preparation**: In preparation for analysis using the Barnes-Hut-SNE algorithm, the data set was log2 transformed. The transcript level for each gene was then normalized across all 74 introgression lines so that the profile had a mean of zero and a standard deviation of one. Normalization of the data allowed for comparison of the relative relationship between each gene expression profile (Bushati et al., 2011).

**Barnes-Hut-SNE**: *t*-SNE or *t*-distributed stochastic neighbor embedding (van der Maaten and Hinton, 2008) is a non-linear dimensionality reduction method, which faithfully maps objects in high dimensional space (H-space) into low dimensional space (V-space). Crowding is avoided through the long-tailed *t*-distribution, which forces non-neighbor clusters farther away from each other in V-space than those clusters actually are in H-space (van der Maaten and Hinton, 2008). The exaggerated separation of non-neighboring clusters improves 2D resolution, allowing identification of novel groupings not readily apparent in other clustering methods. However this method is resource intensive, and with higher dimensionality, the number of genes that can be analyzed is limited. We have used Barnes-Hut-SNE, a newer implementation of *t*-SNE that greatly increases the speed and number of genes that can be analyzed, for the present analysis (van der Maaten, 2013). Barnes-Hut-SNE accomplishes this efficiency through the use of a Vantage Point tree and a variant of the Barnes-Hut algorithm (van der Maaten, 2013). For clustering, 2D maps were generated using a perplexity of 30 and without the initial PCA step from the Barnes-Hut-SNE R implementation (Rtsne package; Krijthe, 2014). Theta was set to 0.3 based on (van der Maaten, 2013) in order to maintain an accurate dimensionality reduction without sacrificing processing speed.

**Clustering for Module Selection**: The DBscan algorithm (Density Based spatial clustering of applications with noise) was used to select modules from the Barnes-Hut-SNE results (fpc package; Hennig, 2014). This algorithm had the advantage of both selecting modules and removing any genes that fell between modules. The scanning range (*epsilon*) and minimum seed points (*minpts*) were selected manually, and used to determine if any one point is a member of a cluster based on physical positioning within the mapping relative to neighboring points. A *minpts* of 25 was used to capture smaller modules on the periphery and an *epsilon* of 2.25 was used to avoid the overlapping of internal and closely spaced modules.

**Plots**: Boxplots were generated from normalized transcript abundance values for each module. The ribbon plot was generated from correlated abundance values from leaf development and photosynthesis related modules. These plots were generated using ggplot form the ggplot2 R Package (Wickham, 2009). The median transcript levels of the genes mapped to a module was calculated for each IL and replicated for all modules. Significant ILs were identified as those with a median transcript level greater than 1 standard deviation from the mean of all genes across all ILs in the module.

### GO Enrichment analysis

Differentially expressed genes at the transcript level for individual ILs and genes with significant eQTL were analyzed for enrichment of Gene Ontology (GO) terms at a 0.05 false discovery rate cutoff (goseq Bioconductor package; Young et al., 2010).

### Promoter enrichment analysis

Promoter enrichment analysis was performed by analyzing the 1000 bp upstream of the ATG translational start site for genes with significant eQTL using 100 motifs represented in the AGRIS AtTFDB (http://arabidopsis.med.ohio-state.edu/AtTFDB). The Biostrings package was used to analyze the abundance of 100 motifs in groups of genes with significant eQTL compared to motif abundance in promoters of all analyzed genes using a Fisher’s exact test (p < 0.05) with either zero or one mismatch (Ichihashi et al., 2014).

### Dissection of eQTL to different stages and time of development at shoot apex

Differentially expressed genes with enriched transcript levels in laser microdissected SAM/P0 vs. P1 samples or hand-dissected samples of the SAM + P0-P4 or P5 sampled over developmental time were obtained from Chitwood et al., 2015. Genes for which a differential expression call could be made (i.e., had enough reads and passed quality filters) were merged with detected eQTL using the merge() function in R (R Core Team, 2015). For bootstrapping, *cis*-and *trans* regulated transcripts were analyzed separately. Merged transcript abundance patterns were randomly permuted (without replacement) using the sample() function against bin identity. For each test, 10,000 permutations were sampled to count the times that a particular transcript expression pattern was assigned to a bin more than the actual count. Resulting frequencies, representing a probability value, were multiple test adjusted using the Benjamini-Hochberg (Benjamini and Hochberg, 1995) method using p.adjust(). Those bins with multiple test adjusted probability values <0.05 were analyzed further using visualizations created with ggplot2 (Wickham, 2009).

### Sequence submission

The quality filtered, barcode-sorted and trimmed short read dataset, which was used to get the normalized read counts and for differential gene expression analysis, was deposited to the NCBI Short Read Archive under accessions SRR1013035 – SRR1013343 (Bioproject accession SRP031491).

## Supplemental files

<u>**Supplemental figures:**</u>

**Supplemental Figure S1. Number of genes in the introgression region for an IL and the number of differentially expressed genes at the transcript level compared to cv. M82**. Strong correlation was observed for differentially expressed genes in *cis* (A), whereas a weak correlation was observed for genes in *trans* (B).

**Supplemental Figure S2. Histograms for differentially expressed genes at the transcript level for the ILs**. Genes with increased or decreased transcript levels for the ILs in *cis* (A), and in *trans* (B).

**Supplemental Figure S3. eQTL and the transcript abundance patterns they regulate**. Each blue bar is a unique introgression. When the transcript abundance pattern of gene 1 is correlated with bin-A, then bin-A contains a *cis*-eQTL. When the transcript abundance pattern of gene 1 is correlated with bin-E then bin-E contains a *trans*-eQTL. When gene 2 has a *cis*-eQTL designated for bin-D and the transcript abundance pattern of gene 2 is also correlated with bin-B, then this secondary correlation is not designated as an eQTL, since these bins share overlapping introgression regions. When the transcript abundance pattern of gene 2 is correlated with bin-B and gene 2 does not have a *cis*-eQTL designated for bin-D, then bin-B is designated as a *trans*-eQTL. All eQTLs for genes that lie in the unassembled portion of the genome (not on any chromosome) cannot be designated as either cis or *trans* and are designated *chromo0*-eQTL.

**Supplemental Figure S4. *Cis*-and *Trans*-eQTL**. Histograms plotting the numbers of significant eQTL mapped to each bin across the 12 chromosomes of *S. lycopersicum* (M82). A. *cis*-eQTLs. B. *trans*-eQTLs.

**Supplemental Figure S5. Frequency and distribution of differentially expressed genes at the transcript level for the IL population at the introgression and the bin level**. (A) Frequency of genes differentially expressed in one or more ILs. (B) Frequency of genes being regulated by one or more BINs/eQTL.

**Supplemental Figure S6. Boxplots of the normalized transcript levels for the three landmark modules**. The relative transcript levels of all genes found in each module for the 74 ILs. The y-axis is the relative transcript level of all eQTL for each IL. (A) Photosynthesis module. (B) Defense, metabolism, and signaling module. (C) Cysteine-type peptidase activity module. Asterisks represent ILs with a median transcript level significantly different from the module mean.

**Supplemental Figure S7. Normalized transcript levels of the leaf development module and genes associated with leaf development within the mapping**. (A) Boxplot of normalized transcript level for the leaf development module. The 108 genes contained in the leaf development module and their relative median transcript level for each IL. Asterisks represent ILs with a median transcript levels significantly different from the module mean. (B) Literature-curated genes related to leaf development (Ichihashi et al., 2014) overlaid on the Leaf Development and Photosynthesis modules. False colored orange, dark blue, and light blue respectively.

**Supplemental Figure S8. eQTL regulation of transcript abundance patterns that correlate with leaf number**.

A, B) Forty-two distinct modules identified by DBscan from the eQTL mapping generated by BH-SNE analysis. Modules enriched for genes with leaf development and photosynthesis GO terms are labeled in blue and green, respectively. Genes with transcript levels correlated with leaf number under simulated sun (A) and shade (B) are indicated by squares with positive correlations in red and negative correlations in yellow.

C, D) Genes with transcript levels correlated with leaf number under simulated sun (C) and shade (D) are shown connected to their respective eQTL with chords. I) The 12 tomato chromosomes in megabases. II) Colored boxes indicate the sizes of each bin. III) Black bars indicate the locations of the genes. IV) Chords connect eQTL to the genes whose transcript levels they regulate. Chords are colored by the chromosome location of the eQTL.

**Supplemental Figure S9. eQTL regulation of transcript abundance patterns that correlate with leaf complexity**.

A, B) Forty-two distinct modules identified by DBscan from the eQTL mapping generated by BH-SNE analysis. Modules enriched for genes with leaf development and photosynthesis GO terms are labeled in blue and green, respectively. Genes with transcript levels correlated with leaf complexity under simulated sun (A) and shade (B) are indicated by squares with positive correlations in red and negative correlations in yellow.

C, D) Genes with transcript levels correlated with leaf complexity under simulated sun (C) and shade (D) are shown connected to their respective eQTL with chords. I) The 12 tomato chromosomes in megabases. II) Colored boxes indicate the sizes of each bin. III) Black bars indicate the locations of the genes. IV) Chords connect eQTL to the genes whose transcript levels they regulate. Chords are colored by the chromosome location of the eQTL.

**Supplemental Figure S10. Distributions of introgressions from *S. pennellii* into *S. lycopersicum* cv. M82**. Map of chromosomes 2, 10, and 11 showing the locations of the introgressions for BIL 033 and 128, as well as the overlapping IL regions, which define the bind (Modified from Chitwood et al., 2013).

**Supplemental Figure S11. Tomato hypocotyl length under sun and shade treatments**. M82 shows a typical shade response with a significantly longer hypocotyl in the shade (Δ of 7 mm). IL 10.1 and BIL 128, which share an overlapping introgression (**Supplemental Fig. S10**), do not significantly respond to the shade treatments. The presence of a response in BIL 033 in combination with the shared introgression with BIL 128 on chromosome 2, indicates that the gene region responsible for the lack of shade response in BIL 128 is located in the introgression on chromosome 10. Bars represent means +/-SE with a minimal of N = 22 for each (ANOVA, F_7,182_ = 44.6, *p* < 0.001). Letters indicate differences at the *p* < 0.05 significance level for Tukey pairwise tests.

<u>**Supplemental tables:**</u>

**Supplemental Table I. Number of differentially expressed (DE) genes at the transcript level in *cis*, *trans*, and the total number of DE genes for the ILs along with number of genes in the introgression region**.

**Supplemental Table II. Correlation coefficients (Spearman’s rho) for significant eQTLs**. The significant eQTL are classified into *trans, cis*, and *Chromo0*, then designated as up (positive slope) or down (negative slope) based on the correlation coefficients.

**Supplemental Table III. GO enrichment and *cis* or *trans* regulation of the 42 identified modules**. All 42 distinct modules are listed with the total number of genes present in each module. The GO enrichment (if one is present) is given for each module, along with whether that module is predominantly *cis* or *trans* regulated. Only nine of the forty-two module show *trans* correlation, which includes the leaf development module.

**Supplemental Table IV. Leaf developmental phenotypes of selected ILs and genetic effects of eQTL (bin) on transcript levels of candidate genes**. Some of the ILs with strong leaf phenotypes are listed along with the associated regulatory QTL (bin) that regulate the transcript levels of candidate leaf-developmental genes.

**Supplemental Table V. Significant correlations between transcript expression patterns and phenotypes**. Bootstrapping analyses correlated transcript expression patterns across the 74 ILs with three phenotypes in under both sun and shade treatments. Genes with significant correlations that also have eQTL are listed.

**<u>Supplemental datasets</u>:**

**Supplemental Dataset 1. List of Differentially Expressed Genes at the transcript level**. List of significant (FDR < 0.05) differentially expressed genes for each Introgression line (IL) compared to cultivated parent *Solanum lycopersicum* cv. M82. For each IL, gene ID, log Fold Change (logFC), log Counts Per Million (logCPM), P-value, False Discovery Rate (FDR) as well as annotation of the differentially expressed genes are presented. The differentially expressed genes for each IL are listed on separate sheets of the file as mentioned on the label of each sheet.

**Supplemental Dataset 2. Transgressive expression of genes at the transcript level among ILs**. Detailed results on transgressive transcript levels of genes for the IL population is presented along with the relevant diagrams and enriched GO-categories for the genes showing transgressive transcript level.

**Supplemental Dataset 3. Genes with Transgressive transcript level**. List of the genes showing transgressive transcript level in the IL population along with details of their transcript abundance and annotation.

**Supplemental Dataset 4. All genes with significant eQTL**. Gene locations are specified by columns: chromosome, bin, binIL.code1-4, begin, end, and sort.order. eQTL bin locations are specified by columns: cor.bin, cor.bin.ILcode1-4, and bin.order. eQTL statistics are in columns: slope.rho.spearman, pvalue.spearman, BHpvalue.spearman, up.down, and cis.trans. AGI and ITAG annotations are listed for the genes in columns V-Y.

**Supplemental Dataset 5. Number of eQTL and genes per bin**. Numbers of *trans*-, *cis*-, and *chromo0*-eQTL are reported for each bin and per gene within each bin.

**Supplemental Dataset 6. Number of eQTL and the bin on which those eQTL reside for each of the landmark modules and Leaf Development module**. The Photosynthesis, Defense, Cysteine-Type Peptidase Activity and Leaf Development modules are listed with the bins on which their eQTL reside. The number eQTL per bin and the percentage of total eQTL within each module are listed.

**Supplemental Dataset 7. Photosynthesis module gene list**. Gene ID of all genes within the Photosynthesis module with description.

**Supplemental Dataset 8. Defense module gene list**. Gene ID of all genes within the Defense, Metabolism and Signaling module with description.

**Supplemental Dataset 9. Cysteine-type module gene list**. Gene ID of all genes within the Cysteine-type Peptidase module with description.

**Supplemental Dataset 10. GO Enrichment for DE Genes**. Enriched GO and GOslim categories for the genes with increased and decreased transcript levels in each IL compared to cultivated parent *Solanum lycopersicum* cv. M82.

**Supplemental Dataset 11. GO Enrichment for eQTL**. Enriched GO and GOslim categories for all the eQTL mapped to a bin, as well as the *cis*-and *trans*-eQTL separately.

**Supplemental Dataset 12. Promoter Motif Enrichment for DE Genes**. Enriched promoter motifs with no mismatch for the up-and down-regulated genes in each IL compared to cultivated parent *Solanum lycopersicum* cv. M82 are shown along with the p-value for significance.

**Supplemental Dataset 13. Promoter Motif Enrichment for *trans*-eQTL**. eQTL are grouped by bin (cor.bin) and promoters were analyzed for both zero and one mismatch with p-values for each. Bins can be ordered using the column, bin.order.

**Supplemental Dataset 14. Differentially expressed leaf Developmental Genes at the transcript level**. Frequency of differentially expressed literature-curated leaf-developmental genes among the ILs. Genes that show differential transcript level in at least one IL are listed. Genes, chromosomes, annotation and the number of ILs showing differential transcript level of the genes are listed

**Supplemental Dataset 15. Curated list of leaf developmental genes with eQTLs**. Gene locations are specified by columns: chromosome, bin, binIL.code1-4, begin, and end. eQTL bin locations are specified by columns: cor.bin, and cor.bin.ILcode1-4. eQTL statistics are in columns: slope.rho.spearman, pvalue.spearman, BHpvalue.spearman, up.down, and cis.trans. AGI annotations for the genes are in columns: Arabidopsis_orthologue and Gene_name.

**Supplemental Dataset 16. Literature-curated plus list of leaf development genes present in the leaf development modules**. Nineteen genes from the literature-curated plus list (Ichihashi et al., 2014) were present within the leaf development module, representing approximately 3% of the curated genes and 11% of the curated genes found in the eQTL data set. The Gene ID, *Arabidopsis* orthologue, and a description for each gene is provided.

**Supplemental Dataset 17. Literature-curated plus list of leaf development genes that are present in all modules**. A total of 175 genes from the curated list plus of leaf developmental genes (Ichihashi et al., 2014) were present in the 5,289 genes with significant eQTL. Matching gene number is the number of genes in the module, which were found in the curated list. The percent of curated genes defines the percentage of matched genes in a module out of the total curated list plus. The percent total in eQTL represents the percentage of matched genes in a module out of the 175 present only in the eQTL. The Photosynthesis and leaf development modules contain the highest proportion of leaf development curated genes.

**Supplemental Dataset 18. Leaf Development module gene list**. Gene IDof all genes within the Leaf Development module with description.

**Supplemental Dataset 19: GO enrichment for bins statistically enriched for transcripts expressed spatio-temporally across tissues**. Enriched GO and GOslim categories for all transcripts expressed spatio-temporally across tissues and with *trans*-eQTL residing in a bin.

**Supplemental Dataset 20: Leaf complexity phenotype of ILs**. Total leaf complexity data for all the ILs under simulated sun and shade is listed.

**Supplemental Dataset 21: Hypocotyl phenotype of ILs**. Hypocotyl length data for all the ILs under simulated sun and shade for two years is listed.

## Acknowledgments

We thank Lauren R. Headland, Jason Kao and Paradee Thammapichai for help in generating plant materials. We also thank Mike Covington for his advice on bioinformatic analyses. We thank the Vincent J. Coates Genomics Sequencing Laboratory at UC Berkeley (supported by NIH S10 Instrumentation Grants S10RR029668 and S10RR027303), and computational resources/cyber infrastructure provided by the iPlant Collaborative (www.iplantcollaborative.org), funded by the National Science Foundation (Grant DBI-0735191).

## Authors' Contributions

DHC, JNM and NRS conceived and designed the experiments. AR, DHC, RK, LC, YI and KZ performed the experiments. AR, JMB and SDR analyzed the data. AR, JMB, SDR, DHC and JNM contributed reagents/materials/analysis tools. AR, JMB, SDR and NRS wrote the paper. All authors read and approved the final manuscript.

## References

AbeM, KatsumataH, KomedaY, TakahashiT (2003) Regulation of shoot epidermal cell differentiation by a pair of homeodomain proteins in Arabidopsis. Development 130: 635–643

AbramoffMD, MagalhaesPJ, RamSJ (2004) Image Processing with ImageJ. Biophotonics International 11: 36–42

BenjaminiY, HochbergY (1995) Controlling the false discovery rate: A practical and powerful approach to multiple testing. J R Stat Soc B 57: 289–300

BremRB, YvertG, ClintonR, KruglyakL (2002) Genetic dissection of transcriptional regulation in budding yeast. Science 296: 752–755

BushatiN, SmithJ, BriscoeJ, WatkinsC (2011) An intuitive graphical visualization technique for the interrogation of transcriptome data. Nucleic Acids Res 39: 7380–7389

CarlesCC, Choffnes-InadaD, RevilleK, LertpiriyapongK, FletcherJC (2005) ULTRAPETALA1 encodes a SAND domain putative transcriptional regulator that controls shoot and floral meristem activity in Arabidopsis. Development 132: 897–911

ChenX, HackettCA, NiksRE, HedleyPE, BoothC, DrukaA, MarcelTC, VelsA, BayerM, MilneI, MorrisJ, RamsayL, MarshallD, CardleL, WaughR (2010) An eQTL analysis of partial resistance to Puccinia hordei in barley. PLoS One 5: e8598

ChitwoodDH, KumarR, HeadlandLR, RanjanA, CovingtonMF, IchihashiY, FulopD, Jimenez-GomezJM, PengJ, MaloofJN, SinhaNR (2013) A quantitative genetic basis for leaf morphology in a set of precisely defined tomato introgression lines. Plant Cell 25: 2465–2481

ChitwoodDH, KumarR, RanjanA, PelletierJM, TownsleyBT, IchihashiY, MartinezCC, ZumsteinK, HaradaJJ, MaloofJN, SinhaNR (2015) Light-Induced Indeterminacy Alters Shade-Avoiding Tomato Leaf Morphology. Plant Physiol 169: 2030–2047

ChitwoodDH, RanjanA, KumarR, IchihashiY, ZumsteinK, HeadlandLR, Ostria-GallardoE, Aguilar-MartinezJA, BushS, CarriedoL, FulopD, MartinezCC, PengJ, MaloofJN, SinhaNR (2014) Resolving distinct genetic regulators of tomato leaf shape within a heteroblastic and ontogenetic context. Plant Cell 26: 3616–3629

ChitwoodDH, SinhaNR (2013) A census of cells in time: quantitative genetics meets developmental biology. Curr Opin Plant Biol 16: 92–99

ClarkRM, WaglerTN, QuijadaP, DoebleyJ (2006) A distant upstream enhancer at the maize domestication gene tb1 has pleiotropic effects on plant and inflorescent architecture. Nat Genet 38: 594–597

CnopsG, Jover-GilS, PetersJL, NeytP, De BlockS, RoblesP, PonceMR, GeratsT, MicolJL, Van LijsebettensM (2004) The rotunda2 mutants identify a role for the LEUNIG gene in vegetative leaf morphogenesis. J Exp Bot 55: 1529–1539

CubillosFA, CousthamV, LoudetO (2012) Lessons from eQTL mapping studies: non-coding regions and their role behind natural phenotypic variation in plants. Curr Opin Plant Biol 15: 192–198

DeCookR, LallS, NettletonD, HowellSH (2006) Genetic regulation of gene expression during shoot development in Arabidopsis. Genetics 172: 1155–1164

Diaz-MendozaM, Velasco-ArroyoB, Gonzalez-MelendiP, MartinezM, DiazI (2014) C1A cysteine protease-cystatin interactions in leaf senescence. J Exp Bot 65: 3825–3833

DrukaA, PotokinaE, LuoZ, JiangN, ChenX, KearseyM, WaughR (2010) Expression quantitative trait loci analysis in plants. Plant Biotechnol J 8: 10–27

EshedY, ZamirD (1995) An introgression line population of Lycopersicon pennellii in the cultivated tomato enables the identification and fine mapping of yield-associated QTL. Genetics 141: 1147–1162

FraryA, NesbittTC, GrandilloS, KnaapE, CongB, LiuJ, MellerJ, ElberR, AlpertKB, TanksleySD (2000) fw2.2: a quantitative trait locus key to the evolution of tomato fruit size. Science 289: 85–88

FridmanE, CarrariF, LiuYS, FernieAR, ZamirD (2004) Zooming in on a quantitative trait for tomato yield using interspecific introgressions. Science 305: 1786–1789

FukaoT, XuK, RonaldPC, Bailey-SerresJ (2006) A variable cluster of ethylene response factor-like genes regulates metabolic and developmental acclimation responses to submergence in rice. Plant Cell 18: 2021–2034

FulopD, RanjanA, OfnerI, CovingtonMF, ChitwoodDH, WestD, IchihashiY, HeadlandL, ZamirD, MaloofJN SinhaNR (2016) A new advanced backcross tomato population enables high resolution leaf QTL mapping and gene identification. bioRxiv 040923; doi: http://dx.doi.org/10.1101/040923

GoffSA, VaughnM, McKayS, LyonsE, StapletonAE, GesslerD, MatasciN, WangL, HanlonM, LenardsA, MuirA, MerchantN, LowryS, MockS, HelmkeM, KubachA, NarroM, HopkinsN, MicklosD, HilgertU, GonzalesM, JordanC, SkidmoreE, DooleyR, CazesJ, McLayR, LuZ, PasternakS, KoesterkeL, PielWH, GreneR, NoutsosC, GendlerK, FengX, TangC, LentM, KimSJ, KvilekvalK, ManjunathBS, TannenV, StamatakisA, SandersonM, WelchSm, CranstonKA, SoltisP, SoltisD, O’MearaB, AneC, BrutnellT, KleibensteinDJ, WhiteJW, Leebens-MackJ, DonoghueMJ, SpaldingEP, VisionTJ, MyersCR, LowenthalD, EnquistBJ, BoyleB, AkogluA, AndrewsG, RamS, WareD, SteinL, StanzioneD (2011) The iPlant Collaborative: Cyberinfrastructure for Plant Biology. Front Plant Sci 2: 34

HammondJP, MayesS, BowenHC, GrahamNS, HaydenRM, LoveCG, SpracklenWP, WangJ, WelhamSJ, WhitePJ, KinggJ, BroadleyMR (2011) Regulatory hotspots are associated with plant gene expression under varying soil phosphorus supply in Brassica rapa. Plant Physiol 156: 1230–1241

HennigC (2014) FPC: Flexible procedures for Clustering. R Package Version: 2.1–9

HollowayB, LiB (2010) Expression QTLs: applications for crop improvement. Molecular Breeding 26: 381–391

HoltanHE, HakeS (2003) Quantitative trait locus analysis of leaf dissection in tomato using Lycopersicon pennellii segmental introgression lines. Genetics 165: 1541–1550

IchihashiY, Aguilar-MartinezJA, FarhiM, ChitwoodDH, KumarR, MillonLV, PengJ, MaloofJN, SinhaNR (2014) Evolutionary developmental transcriptomics reveals a gene network module regulating interspecific diversity in plant leaf shape. Proc Natl Acad Sci U S A 111: E2616–2621

JansenRC, NapJp (2001) Genetical genomics: the added value from segregation. Trends Genet 17: 388–391

KeurentjesJJ, FuJ, TerpstraIR, GarciaJM, van den AckervekenG, SnoeklB, PeetersAJ, VreugdenhilD, KoornneefM, JansenRC (2007) Regulatory network construction in Arabidopsis by using genome-wide gene expression quantitative trait loci. Proc Natl Acad Sci U S A 104: 1708–1713

KliebensteinD (2009) Quantitative genomics: analyzing intraspecific variation using global gene expression polymorphisms or eQTLs. Annu Rev Plant Biol 60: 93–114

KliebensteinDJ, WestMA, van LeeuwenH, KimK, DoergeRW, MichelmoreRW, St ClairDA (2006) Genomic survey of gene expression diversity in Arabidopsis thaliana. Genetics 172: 1179–1189

KoenigD, Jimenez-GomezJM, KimuraS, FulopD, ChitwoodDH, HeadlandLR, KumarR, CovingtonMF, DevisettyUK, TatAV, TohgeT, BolgerA, SchneebergerK, OssowskiS, LanzC, XiongG, Taylor-TeeplesM, BradySM, PaulyM, WeigelD, UsadelB, FernieaR, PengJ, SinhaNR, MaloofJN (2013) Comparative transcriptomics reveals patterns of selection in domesticated and wild tomato. Proc Natl Acad Sci U S A 110: E2655–2662

KrijtheJ (2014) Rtsne: T-distributed Stochastic Neighbor Embedding using Barnes-Hut implementation. R Package Version: 0.9

KroymannJ, DonnerhackeS, SchnabelrauchD, Mitchell-OldsT (2003) Evolutionary dynamics of an Arabidopsis insect resistance quantitative trait locus. Proc Natl Acad Sci U S A 100 Suppl 2: 14587–14592

KumarR, IchihashiY, KimuraS, ChitwoodDH, HeadlandLR, PengJ, MaloofJN, SinhaNR (2012) A High-Throughput Method for Illumina RNA-Seq Library Preparation. Front Plant Sci 3: 202

LiH, DurbinR (2009) Fast and accurate short read alignment with BurrowsWheeler transform. Bioinformatics 25: 1754–1760

LiuYS, ZamirD (1999) Second generation L. pennellii introgression lines and the concept of bin mapping. Tomato Genet. Coop. Rep. 49: 26–30

LongairMH, BakerDA, ArmstrongJD (2011) Simple Neurite Tracer: open source software for reconstruction, visualization and analysis of neuronal processes. Bioinformatics 27: 2453–2454

MoyleLC (2008) Ecological and evolutionary genomics in the wild tomatoes (Solanum sect. Lycopersicon). Evolution 62: 2995–3013

MuirCD, PeaseJB, MoyleLC (2014) Quantitative genetic analysis indicates natural selection on leaf phenotypes across wild tomato species (Solanum sect. Lycopersicon; Solanaceae). Genetics 198: 1629–1643

MullerNA, WijnenCL, SrinivasanA, RyngajlloM, OfnerI, LinT, RanjanA, WestD, MaloofJN, SinhaNR, HuangS, ZamirD, Jimenez-GomezJM (2016) Domestication selected for deceleration of the circadian clock in cultivated tomato. Nat Genet 48: 89–93

R Development CoreTeam (2015) R: A Language and Environment for Statistical Computing. (Vienna, Austria: R Foundation for Statistical Computing)

RanjanA, IchihashiY, SinhaNR (2012) The tomato genome: implications for plant breeding, genomics and evolution. Genome Biol 13: 167

RobertsA, PachterL (2013) Streaming fragment assignment for real-time analysis of sequencing experiments. Nat Methods 10: 71–73

RobinsonMD, OshlackA (2010) A scaling normalization method for differential expression analysis of RNA-seq data. Genome Biol 11: R25

SchadtEE, MonksSA, DrakeTA, LusisAJ, CheN, ColinayoV, RuffTG, MilliganSB, LambJR, CavetG, LinsleyPS, MaoM, StoughtonRB, FriendSH (2003) Genetics of gene expression surveyed in maize, mouse and man. Nature 422: 297–302

SharlachM, DahlbeckD, LiuL, ChiuJ, Jimenez-GomezJM, KimuraS, KoenigD, MaloofJN, SinhaN, MinsavageGV, JonesJB, StallRE, StaskawiczBJ (2013) Fine genetic mapping of RXopJ4, a bacterial spot disease resistance locus from Solanum pennellii LA716. Theor Appl Genet 126: 601–609

SvistoonoffS, CreffA, ReymondM, Sigoillot-ClaudeC, RicaudL, BlanchetA, NussaumeL, DesnosT (2007) Root tip contact with low-phosphate media reprograms plant root architecture. Nat Genet 39: 792–796

van der MaatenL (2013) Barnes-Hut-SNE. arXiv:1301.3342[cs.LG]

van der MaatenL, HintonG (2008) Visualizing Data using t-SNE. Journal of Machine Learning Research 9: 2579–2605

WernerJD, BorevitzJO, WarthmannN, TrainerGT, EckerJR, ChoryJ, WeigelD (2005) Quantitative trait locus mapping and DNA array hybridization identify an FLM deletion as a cause for natural flowering-time variation. Proc Natl Acad Sci U S A 102: 2460–2465

WestMA, KimK, KliebensteinDJ, van LeeuwenH, MichelmoreRW, DoergeRW, St ClairDA (2007) Global eQTL mapping reveals the complex genetic architecture of transcript-level variation in Arabidopsis. Genetics 175: 1441–1450

WickhamH (2009) ggplot2: elegant graphics for data analysis. Springer New York

YoungMD, WakefieldMJ, SmythGK, OshlackA (2010) Gene ontology analysis for RNA-seq: accounting for selection bias. Genome Biol 11: R14

YuD, ChenC, ChenZ (2001) Evidence for an important role of WRKY DNA binding proteins in the regulation of NPR1 gene expression. Plant Cell 13: 1527–1540

ZhangL, FetchT, NirmalaJ, SchmiererD, BrueggemanR, SteffensonB, KleinhofsA (2006) Rpr1, a gene required for Rpg1-dependent resistance to stem rust in barley. Theor Appl Genet 113: 847–855

